# Influence of *Oenococcus oeni* and *Brettanomyces bruxellensis* on Aged Wine Microbial Taxonomic and Functional Profiles

**DOI:** 10.1101/216390

**Authors:** Marie Lisandra Zepeda-Mendoza, Nathalia Kruse Edwards, Mikkel Gulmann Madsen, Martin Abel-Kistrup, Lara Puetz, Thomas Sicheritz-Ponten, Jan H. Swiegers

## Abstract

In the wine making process, the interactions between lactic acid bacteria (LAB), yeast and other wine microflora have an impact on the wine quality. In this study, we investigate the influence of the LAB *Oenococcus oeni* and the spoilage yeast *Brettanomyces bruxellensis* on the microbial community of a Cabernet Sauvignon wine. We generated metagenomic datasets from inoculations of three strains of *B. bruxellensis*, in combination with two *O. oeni* strains, one with and one without cinnamoyl esterase activity. This esterase activity releases hydroxycinnamic acids (HCAs) that can subsequently be processed by some *B. bruxellensis* strains able to generate off-flavor compounds. We evaluated the influence of the *O. oeni* and *B. bruxellensis* on the microbial taxonomic and functional potential profile, particularly regarding off-flavor formation due to HCAs. We found that the effect on the microbial profiles depends on *i*) the *O. oeni* and *B. bruxellensis* strains being combined and *ii*) the abundance they reach in the final wine, which depends on certain unidentified conditions. We confirmed that the potential of *B. bruxellensis* to produce off-flavor compounds from HCAs depends on the strain. Interestingly, the samples without microbial inoculants also had this potential, suggesting that native grape microbiota could also influence the levels of HCA. We also found that the presence of *B. bruxellensis* does not interfere with the malolactic fermentation of the evaluated *O. oeni* strains, which leads to a less acidic taste. We show that metagenomic approaches can help uncover the complex wine microbial community traits, such as flavor, impacted by the simultaneous presence of *O. oeni* and *B. bruxellensis*.

## Introduction

### Wine-making and wine microbial interactions

The study of the wine microbial ecosystem has mostly been focused on the two key fermentation players, *Saccharomyces cerevisiae*, which performs the alcoholic fermentation (AF), and *Oenococcus oeni,* which performs the malolactic fermentation (MLF), although other genus and species can also affect the wine characteristics. MLF is in fact a decarboxylation process where dicarboxylic L-malic acid (malate) is converted to monocarboxylic L-Lactic acid (lactate) and CO^2^, which can result in a rise in pH. Thus, MLF softens the acid structure of the wine, as L-lactic acid is perceived as less acidic than L-malic acid. MLF usually starts spontaneously about 1-3 weeks after completion of AF and lasts 2-12 weeks. Furthermore, some carbohydrates are metabolized during MLF and there is release of phenolic acids and synthesis of esters, among other reactions, which are important for the wine flavour profile (Lonvaud-Funel n.d.). LAB naturally present in wine, or commercial strains isolated from wine, are normally used for MLF, with *O. oeni* being the preferable species due to its ethanol and acid tolerance and flavour profile (reviewed by Liu 2002 (Liu 2002)).

In the process of fermented food and beverage making, the starting ingredients can have associated indigenous microbial communities (Leff et al. 2013), which might vary depending on the source and could have an impact on the final product characteristics. This has been investigated in wine, where the grape microbiota is influenced by cultivar, vintage, and climate (Bokulich et al. 2014). Besides yeasts, such as *Saccharomyces*, and filamentous fungi, such as *Aspergillus* and *Penicillium*, a large bacterial diversity has been observed on grapes and must. The bacterial community is formed mainly by Proteobacteria, including acetic acid bacteria, and Firmicutes, including lactic acid bacteria (LAB) (reviewed by Barata et al. (2012) (Barata et al. 2012)). Some bacteria are plant or environmental microbes, while others have the physiological characteristics to allow them to grow on the harsh oenological conditions (Pina et al. 2004) (low nutrients, high acidity, ethanol concentrations of up to 15% v/v), thus being able to form part of the wine microbiome (Barata et al. 2012). The microbial interactions, as well as their succession dynamics through the wine making process affect the hygienic and organoleptic properties of the final wine product (Sieuwerts et al. 2008). For example, *Botrytis cinerea* influences the microbial taxonomic profile through release of nutrients (Barata et al. 2008); one of those affected yeasts is the genus *Metschnikowia*, which can in turn prevent the growth of other fungi and bacteria by sequestering iron (Sipiczki 2006). Microbial interactions are also known to be highly strain specific. For example, a strain of *S. cerevisiae* has been shown to produce antimicrobial peptides under oenological conditions, which can inhibit growth of *Brettanomyces bruxellensis* (Branco et al. 2014). In the wine industry, *B. bruxellensis* is a spoilage yeast difficult to get rid of, mostly present in barrel aged wines (reviewed by Suarez *et al.* 2006 (Suarez et al. 2007)).

### Hydroxycinnamic acids

A potential source of off-flavour compounds is hydroxycinnamic acids (HCAs). HCAs are organic acids, naturally present in grapes and wines, however, they are usually found as tartaric acid bound esters in grapes and wine. The content depends on the grape variety and growth conditions (Nagel & Wulf 1979). This family of organic acids has been studied in wine and some food systems for its properties, such as colour stabilizing (Hernández et al. 2006), antioxidant (Bouzanquet et al. 2012), radical scavenging (Kikuzaki et al. 2002), and antimicrobial activity against some yeast and bacteria (Ou & Kwok 2004). However, the full effect of HCAs in food and in wine is not yet fully understood. Some LAB strains of Oenococcus and other Lactobacillus have been shown poses a cinnamoyl esterase activity, which releases HCAs from their bound form (Cabrita et al. 2008). It has been shown that the cinnamoyl esterase can also be present in different fungi (Rumbold et al. 2003). Furthermore, HCAs can also be released by chemical hydrolysis due to the acidity of the wine in a slow process that gradually continues through the winemaking and storage (Hixson et al. 2012). *B. bruxellensis* does not have the capability of producing free HCAs, however some strains possess a decarboxylase and a vinyl reductase activity, which can convert them into the off-flavour volatile phenols 4-ethylphenol and 4-ethylguaiacol, which confer the “Brettanomyces aroma” (Hixson et al. 2012). An increase in free HCAs could hereby potentially increase the risk of spoilage by a *B. bruxellensis* strain with both activities (reviewed by Kheir et al. 2013 (Kheir et al. 2013)). Importantly, Madsen *et al.* (2016) (Madsen et al. 2016) showed that the concentration of volatile phenols depends more on the strain of Brettanomyces than on the HCA esterase activity of *O. oeni*. Thus, the strain of *B. bruxellensis* is key in determining the volatile phenol concentration. Determination of the effect of a specifically selected starter or mixed-starter culture of yeast and bacteria on the wine profile cannot be effectively done without also characterizing the entire microbial community (Liu et al. 2017). Furthermore, although factors affecting yeast biodiversity have been widely documented, less characterization has been performed on the factors that influence the bacterial population. While several studies have characterized the microbiota on the grape surface and must, there is scarcity in the characterization of the microbial community in the final wine product. “Omics” methodologies in the food sciences, in particular in fermented goods, have been applied for the deeper and broader analysis of the microbial system relevant to both the fermentation process and the characteristics of the final product (De Filippis et al. 2017). In this study, we undertook a metagenomic approach to characterize the impact of the inoculations of two different strains of *O. oeni* (with and without the esterase activity) and three *B. bruxellensis* strains, alone and in combination, on the wine microbial community six months post-inoculation in a Cabernet Sauvignon wine. We furthermore characterized the MLF activity in the inoculations by measuring the abundance of malic acid.

## Materials and Methods

### Wine inoculation

Destemmed grape must from Cabernet Sauvignon before AF was imported from Bulgaria by Chr. Hansen A/S. *S. cerevisiae* strain NI6 was used for AF in 50 L tanks. The wine was pressed through filter cloths to remove grape seeds and skin. After mixing the wine by stirring, the wine was decanted into 5 L containers and stored at 5 °C. The wine was measured on an Oenofoss and had an alcohol percent of 12.6 %, 0.0 g/L glucose and 0.0 g/L fructose. The sulphite level was measured with a Megazyme kit and found to be 25 ppm. The pH was measured during MLF and remained at 3.5.

Pure cultures of *B. bruxellensis* were stored on YGC agar to ensure viability of the yeasts throughout the experiment. YGP broth was made with 5.0 g yeast extract, 10.0 g peptone and 11.0 g glucose monohydrate and milli-q water added until 1 L total volume in a conical flask. The mixture was dissolved using a magnetic stirrer. The pH was adjusted to 5.6 ±0.2 with 1 M HCL and 1 M NaOH and hereafter autoclaved at 121 °C for 15 minutes. The broth was kept in a refrigerator at 5 °C until use. CBS 73 had been frozen in glycol and was rejuvenated in YGP broth for 48 hours at 25 °C before being inoculated into a new YGP broth, grown for 72 hours at 25 °C and then plated on YGC agar. The two other strains were directly inoculated in YGP broth and then plated on YGC agar. All strains were incubated at 25 °C. A sample from each flask was counted after 72 hours on a hemocytometer (see 3.2) where the strains reached a level of approx. 10^6^-10^7^ CFU/mL. CBS 73 (Brett_A), CBS 2336 (Brett_B) and CBS 2499 (Brett_C) and were continually grown for 72 hours at 25 °C.

Two different strains of *O. oeni* were provided by Chr. Hansen A/S and kept frozen at -18 °C. One of the *O. oeni* strains is cinnamoyl esterase negative (from here on referred to as OEN), while the other is cinnamoyl esterase positive (from here on referred to as OEP), which can hereby potentially liberate HCAs. 1.2 g frozen culture made for direct inoculation was dissolved in 200 mL sterilized peptone water and 5 mL was added to 1 L of wine corresponding to an inoculation level of approx. 106 CFU/mL. The cell concentrations in the YGP broth right before inoculation were determined using a hemocytometer where cells were counted in the microscope.

Wine was collected from Chr. Hansen A/S in the morning of day 0 of MLF. Wine was poured into 20 autoclaved 1 L blue-cap bottles and 4 autoclaved 5 L blue-cap bottles. Control wines were put aside in a tempered room (20 °C). To minimize cross-contamination, the wines for MLF were then inoculated with OEP and OEN and two wines of each (MLF control) were put aside in a tempered room. The rest of the OEP wines were then inoculated with either Brett_A, Brett_B or Brett_C, one at a time. Afterwards, the wines with OEN were inoculated also with the respective *B. bruxellensis* strains. Control wines containing only *B. bruxellensis* were finally inoculated one strain at a time. *B. bruxellensis* was inoculated at a concentration of approx. 5 × 10^2^ by pipetting.

### Measurement of MLF

An enzyme test-kit was used to measure the malic acid content. Samples were taken on days 0, 4, 7, 10, 14 and 114 from every bottle and frozen for later analysis using a malic acid enzyme test-kit (R-Biopharm, Germany) and absorbance measurements on a spectrophotometer. A cuvette with 1.00 cm light path was used at wavelength 340 nm at 20-25 °C. The cuvettes were prepared according to the kit instructions. We calculated the malic acid concentration using the absorbance values.

### DNA extraction and sequencing

The wines were sampled on day 114 after inoculation. The bottles of wine were gently swirled before sampling to ensure proper mixing of the wine. Sterile B. Braun omnifix syringes without needles were used to take samples from the bottles with a minimum of oxygen intake, although it could not be entirely avoided. The samples were frozen at -60 °C until analysis.

For DNA isolation, cells were pelleted from 50 mL of wine centrifuged at 4500 g for 10 minutes and subsequently washed three times with 10 mL of 4°C phosphate buffered saline (PBS). The pellet was mixed with G2-DNA enhancer (Ampliqon, Odense, Denmark) in 2 ml tubes and incubated at RT for 5 min. Subsequently, 1 mL of lysis buffer (20 mM Tris-HCl-pH 8.0, 2 mM EDTA and 40mg/ml lysozyme) was added to the tube and incubated at 37°C for one hour. An additional 1 mL of CTAB/PVP lysis buffer was added to the lysate and incubated at 65°C for one hour. DNA was purified from 1 mL of lysate with an equal volume of phenol-chloroform-isoamyl alcohol mixture 49.5: 49.5: 1 and the upper aqueous layer was further purified with a MinElute PCR Purification kit and the QIAvac 24 plus (Qiagen, Hilden, Germany), according to the manufacturer’s instructions, and finally eluted in 100 ul DNase-free H_2_O.

Prior to library building, genomic DNA was fragmented to an average length of ~400 bp using the Bioruptor^®^ XL (Diagenode, Inc.), with the profile of 20 cycles of 15 s of sonication and 90 s of rest. Sheared DNA was converted to Illumina compatible libraries using NEBNext library kit E6070L (New England Biolabs) and blunt-ended library adapters described by Meyer and Kircher (2010) (Meyer & Kircher 2010). The libraries were amplified in 100-μL reactions, with each reaction containing 20 μL of template DNA, 10 U AmpliTaq Gold polymerase (Applied Biosystems, Foster City, CA), 1× AmpliTaq Gold buffer, 2.5 mM MgCl2, 0.2 mM of each dNTP, 0.2 uM IS4 forward primer and 0.2 uM reverse primer with sample specific 6 bp index. The PCR conditions were 12 minutes at 95°C to denature DNA and activate the polymerase, 14 cycles of 95°C for 20 seconds, 60°C annealing for 30 seconds, and 72°C extension for 40 seconds, and a final extension of 72°C extension for 5 minutes. Following amplification, libraries were purified with Agencourt AMPure XP (Beckman Coulter, Inc) bead purification, following manufacturer’s protocol, and eluted in 50 uL of EB buffer (Qiagen, Hilden, Germany). The quality and quantity of the libraries were measured on the Bioanalyzer 2100 (Agilent technologies, Santa Clara, United States), and the libraries were pooled at equimolar concentration. Sequencing was performed on the Illumina HiSeq 2500 in PE100 mode following the manufacturer’s instructions.

### Metagenomic taxonomic profiling

The reads were first cleaned with cutadapt (Martin 2011) to remove adapter sequences and low quality bases (min quality= 33, 3’-end minimum quality= 30, minimum length= 30). In order to evaluate the inoculation efficiency of the *O. oeni* and the *B. bruxellensis* strains, we mapped with bwa v0.7.10 (Li & Durbin 2009) the cleaned reads against the genomes of the *O. oeni* strains (in house genomic sequences) and the published genome of *B. bruxellensis* (CBS 2499 v2.0) and calculated the depth and breadth of coverage using samtools v1.3.1 and bedtools v.2.26. The coverage statistics of *B. bruxellensis* were calculated excluding the scaffolds AHMD01000878.1, AHMD01000885.1 and AHMD01000879.1, which contain rDNA tandem repeats, which we found to artificially inflate the coverage due to mapping of reads likely deriving from other yeasts.

In order to characterize the microbial profiles of the inoculations, we used MGmapper (Petersen et al. 2017) to first map the reads against the *phi* genome. The non-mapping reads were then used to map against the next databases extracted from NCBI (2016/09/20) in “best mode”: human, plant, vertebrates, invertebrates, virus, fungi, protozoa, plasmid, and bacteria. The number of mapping reads, coverage and depth were calculated, and the hits were annotated from the superkingdom to the species taxonomic level. The identifications were filtered by taking into account the next parameters: minimum abundance of 0.01%, minimum ratio of unique mapping reads and total mapping reads of 0.005, maximum edit distance of 0.01, and minimum of 10 mapping reads.

### Metagenomic taxonomic comparison

In order to compare the microbial populations of the different inoculation samples we first built a matrix with the number of reads mapping to the filtered identifications from all the samples and normalized the counts by percentage of abundance. We used this matrix to *i*) identify the core microbiomes of each type of inoculation, *ii*) the diversity distance, *iii*) to perform principal component analysis (PCA), *iv*) differential abundance, and *v*) abundance correlation analyses using R. The comparative analyses were performed excluding the out-layer samples with the highest and lowest depth of sequencing (OEN_B_18 and OEN_23, respectively).

The taxonomic cores were obtained by identifying the microbes present in all the replicates of each inoculation type. We calculated the microbial diversity distance between and within the inoculation types using the R package vegan using the Bray, Jaccard, and Euclidean distances and clustered them with the ward.D and average methods. The differential abundance was performed using Fisher test with alternative hypotheses greater and less. We built the contingency tables using the mean of the technical replicates of the inoculation types and performed the next comparisons: *i*) all the inoculation types versus the controls, *ii*) the combinations of OEP and the three *B. bruxellensis* strains versus OEP, *iii*) the combinations of OEN and the three *B. bruxellensis* strains versus OEN, *iv*) OEP versus OEN, *v*) the combinations of each *B. bruxellensis* strain and the two *O. oeni* strains versus the given *B. bruxellensis* strain. The p values (*P*) were adjusted by the false discovery rate (FDR) and the significant comparisons were those with FDR ≤ 0.05.

The abundance correlations were performed with the R function cor.test using the Spearman method. We removed from the normalized count matrix those identifications present in less than 10 samples. We defined two types of significant correlations (*P* < 0.05 and rho < -0.4 or rho > 0.7): a) Unaffected correlations: the ones identified when comparing all the samples and when comparing without each of the inoculation types. b) Affected correlations: the ones that were identified only when removing one of the inoculation types. In order to identify the top 5% abundant taxa in each sample, we normalized by depth of coverage. We also identified which top abundant species were present in all the technical replicates of each inoculation type.

### Metagenomic functional potential profiling

The presence of HCA decarboxylase gene (*HcD*) in the used *B. bruxellensis* strains was confirmed using lastz (Harris 2007) to identify the genomic region of the *B. bruxellensis* sequence used as reference containing the sequence of *HcD* transcript id HQ693758.1 and using bedtools to extract the coverage of the mappings of the samples OEP_A_10, OEP_B_11, and OEP_C_13 (the ones with the highest coverage of each *B. bruxellensis* strain).

#### Nr gene set catalogue

The cleaned reads were *de novo* assembled using IDBA-UD v1.1.1 (Peng et al. 2012) using the pre-correction parameter in order to account for the uneven sequencing depths. Genes were then predicted on the assemblies with prodigal v2.6.2 (Hyatt et al. 2010) using the meta mode. Afterwards, the predicted genes of each sample were clustered using vsearch v2.1.2 (Edgar 2010) with an identity threshold of 95% and a minimum sequence length of 20. The centroid sequence of each cluster was kept as the representative sequence to form a non-redundant (nr) gene set.

Afterwards, the nr gene sets were pooled and clustered using usearch with the same parameters to generate the final nr gene set catalogue used for the functional potential comparative analyses.

#### Comparative analyses

The reads of each sample were mapped against the nr gene set catalogue using bwa mem (Li & Durbin 2009) to then obtain the coverage of each gene using samtools and bedtools. The coverage was used to build an abundance matrix. We then performed principal component analysis (PCA) on the normalized matrix using the function prcomp from R v3.2.0 with scaling. Given that sample OEN_23 was identified as an extreme outlayer and that the *O. oeni* inoculation did not succeed in this replicate, it was removed from the subsequent comparative functional potential analyses. As another method to evaluate the variation between and within the inoculation types, we calculated the Bray, Jaccard and Euclidean distances with the R package vegan using the abundance matrix and the values were clustered using the average and the ward.D methods.

Next, we assigned a KEGG orthology (KO) to the predicted nr genes using blastx with e-value 0.000001 against the KEGG database. The blast hits were filtered by a minimum bit score of 50 and minimum of 30% identity. A new abundance matrix was built for the genes with a KO identification. Subsequently, we identified KOs in differential abundance, i.e. statistically significant less or more abundant in a given inoculation type when compared to another inoculation type. To this end, we performed in R a Fisher test with the alternative hypothesis of greater and less and corrected *P* using FDR. A contingency table for the Fisher tests using the mean of the replicates was made for comparing each of the inoculation types against the control, the OEP_A/B/C against OEP, OEN_A/B/C against OEN, OEN against OEP, and each of the *O. oeni*_A/B/C against the corresponding *B. bruxellensis* strain. We kept as statistically significant those with a FDR ≤ 5%.

The differentially abundant KOs were then grouped by the pathways they are part of as annotated in KEGG and a two-sided Fisher test was performed as previously described for the KOs using as counts the number of differentially abundant KOs belonging to that pathway. We kept as significant those with *P* ≤ 0.05. Then, we identified those differentially abundant pathways present in a minimum of 4 samples, those present only in the comparisons against the control, the OEP_A/B/C combinations against OEP, the OEN_A/B/C combinations against OEN, OEP against OEN, and the two *O. oeni* strains combined with each of the *B. bruxellensis* strains against the respective *B. bruxellensis* strain.

Furthermore, we identified the functional cores of the different inoculation types, defined as those nr genes present only in anyone of the replicates of each of the inoculation types. We also identified the most abundant genes in each of the samples as those within the top 5% genes with highest counts and those present in the two replicates as top abundant were identified.

## Results

### Taxonomic profiling

A total of 534,135,264 sequencing reads were produced from all the samples (min= 1,899,372, max= 162,426,646, average= 23,223,272.35), from where we obtained 523,937,023 cleaned reads (min=1,831,535, max=159,467,366, average=22,779,870.57) (Table 1, Supplemental File 1). Although two replicates were produced for each inoculation type and control, it was not possible to extract DNA from one of the replicates of the inoculation with OEP and Brett_C. Regarding the inoculation with only OEN, one of the replicates (OEN_23) has the lowest number of reads (1,899,372). Thus, it was removed from the taxonomic and functional comparisons, as it would not capture the low abundant microbes identified by the other samples and it represented an extreme out layer in the evaluation of the functional potential profile with a PCA of all the samples due to its low sequencing depth (Supplemental Figure 1).

**Table 1.** Statistics of the sequencing and the mapping to the genomes of the inoculants.

| Sample | Cleaned reads | % Unmapped reads | *O. oeni* * coverage | *B. bruxellensis* ** coverage |
| --- | --- | --- | --- | --- |
| Control_1 | 3,317,232 | 21 | 2.75 | 0.35 |
| Control_2 | 5,195,056 | 15.23 | 4.45 | 0.88 |
| Brett_A_3 | 4,107,740 | 8.54 | 1.15 | 26.3 |
| Brett_A_4 | 6,695,169 | 8.41 | 18.1 | 38.7 |
| Brett_B_5 | 2,761,092 | 2.08 | 70.65 | 0.4 |
| Brett_B_6 | 10,993,754 | 2.04 | 420 | 0.45 |
| Brett_C_7 | 13,827,278 | 3.25 | 399.8 | 23.27 |
| Brett_C_8 | 2,054,860 | 4.35 | 14.35 | 12.97 |
| OEP_A_9 | 8,067,295 | 7.9 | 43.85 | 27.2 |
| OEP_A_10 | 65,488,400 | 5.54 | 769.5 | 66.26 |
| OEP_B_11 | 59,758,655 | 1.91 | 2369.1 | 7.9 |
| OEP_B_12 | 19,900,342 | 1.89 | 216.25 | 3.37 |
| OEP_C_13 | 39,001,458 | 3.59 | 945.75 | 105.4 |
| OEN_A_15 | 43,950,984 | 4.27 | 1333 | 21 |
| OEN_A_16 | 6,728,890 | 10.18 | 4.3 | 44.6 |
| OEN_B_17 | 37,224,934 | 2.12 | 1280.85 | 2.12 |
| OEN_B_18 | 159,467,366 | 2.08 | 4621.2 | 13 |
| OEN_C_19 | 6,443,410 | 6.41 | 1.25 | 37.4 |
| OEN_C_20 | 6,458,253 | 6.19 | 110.5 | 26.3 |
| OEP_21 | 65,477,885 | 1.17 | 5.8 | 1.29 |
| OEP_22 | 6,451,292 | 9.15 | 8.785 | 1.23 |
| OEN_23 | 1,831,535 | 51 | 0.635 | 0.13 |
| OEN_24 | 7,664,243 | 3.6 | 47.25 | 0.9 |

\*Reported mapping statistics for *O. oeni* derive from the average of the mapping versus the genomes of OEP and OEN. \*\*Reported mapping statistics for *B. bruxellensis* derive from mapping versus the publicly available genome of Brett\_C strain.

The sample OEN_B_18 has the highest number of reads (162,426,646), thus it was removed from the taxonomic comparisons, as it would bias for the identification of the very low abundant species that the other samples would not capture. However, it was not removed from the functional comparisons. Thus, for the taxonomic comparisons, the total number of species level identifications pooling all the samples for the bacterial database was 918 (899 when removing the two out layer samples), 117 plasmids (109 without out layer samples), 18 archaea, 11 viruses, 332 fungi (328 without out layer samples), and 96 protozoa.

### Inoculants abundance and functional evaluation

We evaluated the abundance of *O. oeni* and *B. bruxellensis* in the wines 6 months after inoculation (Table 1). The genome of *O. oeni* is covered at medium and high coverage in the two Brett_B inoculations (70.65x and 420x). Also, one of the replicates of Brett_C has the genome of *O. oeni* at high coverage (399.8x), while the other is present in low abundance (14.3x). From the OEP_A replicates, one has the genome of *O. oeni* covered at medium coverage (43.85x) with the *B. bruxellensis* strain at abundance similar to the other samples, while the other (sample OEP_A_10) has *O. oeni* in high abundance (769.5x) and is also the sample with the second most abundant *B. bruxellensis* abundance (66.26x), even higher than the samples were *B. bruxellensis* was inoculated alone. And OEP_C_13, which does not have a replicate, has the highest *B. bruxellensis* coverage (105.4x). *B. bruxellensis* was identified among the top abundant fungi only in OEP_A_10 and OEP_C_13.

To test whether the identified *O. oeni* bacteria in the wines not inoculated with it derive from the native grape flora, we evaluated the amount of MLF in the samples. We identified malic acid in the samples inoculated with OEN and OEP at day 114, but no malic acid in those not inoculated with our *O. oeni* strains (Figure 1).

**Figure 1.**
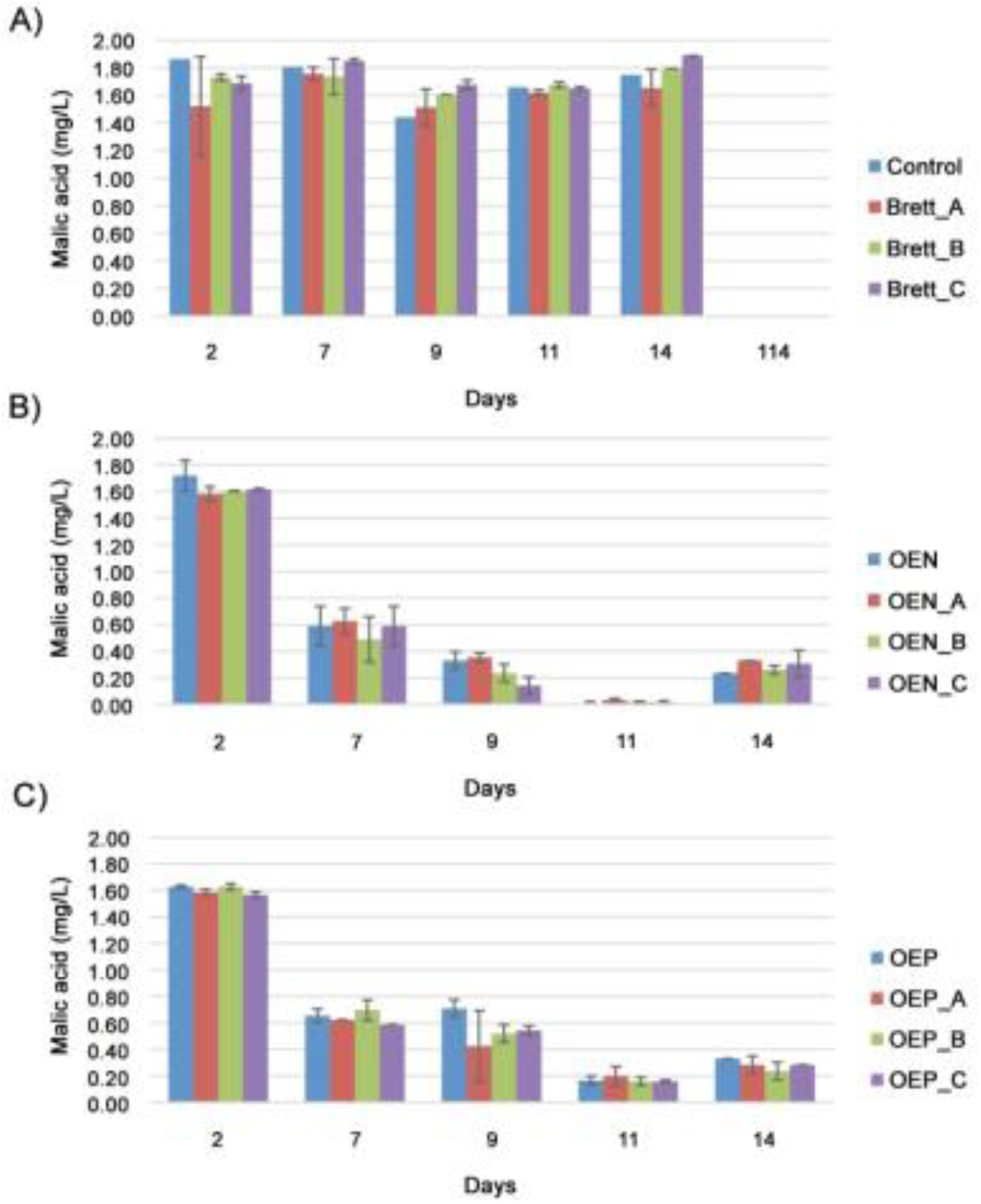
Concentration of malic acid (g/L) over time (days). Malic acid concentrations are average values of duplicates. Error bars are standard deviations. A) Malic acid concentration on inoculations with *B. bruxellensis* strain A, B or C. The concentrations on day 114 were 0 mg/L malic acid in all inoculations, suggesting any identified *O. oeni* in those samples are not able to perform MFL, and are thus likely derived from the grape flora. B) Wines inoculated with OEN and *B. bruxellensis* strain A, B or C. C) Wines inoculated with OEP and *B. bruxellensis* strain A, B or C.

### Bacterial identifications

In the PCA of the identified bacterial species (Figure 2A), most samples cluster tightly together with the controls, with the most variable samples being from the Brett_C inoculation and the Brett_A combined with both *O. oeni* strains. Brett_C_8 has the least number of bacterial identifications (15), however it had much less depth of sequencing than its pair (<25%), but similar sequencing to other samples with more identifications and similar number of identifications to other sample with double sequencing depth (Brett_A_3). Similarly, other inoculation types have a total number of identifications uncorrelated to their depth of sequencing. For example, OEP_A_10 has the highest number of identifications, and the pair has ~1/6 of its sequencing depth (the sample with second highest sequencing). However, it has similar number than OEN_A_15, which has less sequencing than it. Also, the other samples with top highest sequencing have similar number of identifications to the control samples (which have mean depth of sequencing).

**Figure 2.**
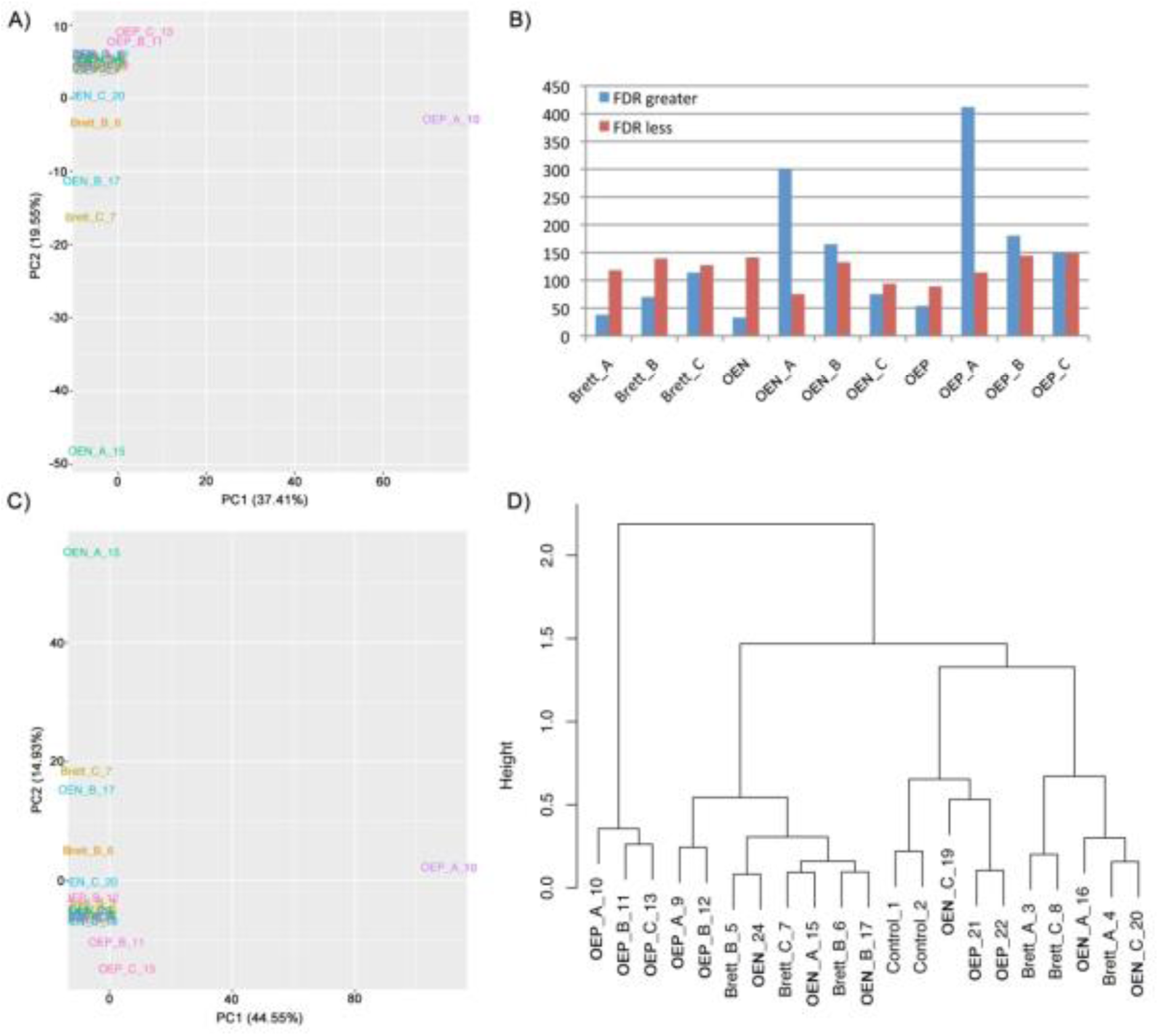
Bacterial profiling. A) PCA of the bacterial taxa at species level. B) Differentially abundant bacterial species by inoculation type compared to the control samples. FDR greater denotes the bacteria that was in statistically significant higher abundance, and FDR less are those in statistically significant less abundant. C) Cladogram of ward.D clustered Bray distances of all the microbial taxa from the samples. D) PCA of the abundance of all the identified microbial taxa.

### Bacterial differential abundance

Regarding the differentially abundant identifications, OEN is the inoculation type with the lowest number of bacteria present in higher abundance compared to the controls (33, while the mean is 144.6 and median 144), Brett_A was the second, with 38 taxa, and OEP is the third with 54. In regards to the number of species in less abundance compared to the controls, the inoculations with only Brett_A have around the average (118, average= 120.1), while OEN has 141 and OEP_C has the maximum (148) (Figure 2B). Interestingly, while Brett_A alone is the one changing the least the bacterial community, it is also the one that changes it the most when inoculated together with *O. oeni* (both OEP and OEN). Comparing the patterns of the number of higher and less abundant species of all the inoculation types compared to the control, we observed that all the inoculation types have similar amount of less abundant bacteria, with the largest difference being only in the number of higher abundant bacteria in OEN_A and OEP_A. OEP_A had the highest number of species present in higher abundance compared to the control (412, while the mean is 144.6 and the median is 144), and OEN_A had the second highest number of species present in higher abundance compared to the control (300). OEN was the one with the lowest number of bacteria present in higher abundance compared to the control (33). Also, compared to Brett_A, both OEN_A and OEP_A have low numbers of differentially less abundant bacteria (24 and 36, respectively). When compared to OEP, OEN has 37 species in differentially higher abundance, and 142 in less abundance.

In evaluating the taxonomic core of the inoculations, we found that the core of the OEN_A inoculation type is the largest (Figure 3A). We also found that the number of species in the OEP_A inoculation type core is not the highest (Figure 3A) while the distance of OEP_A to the control is the largest (Figure 2CD).

**Figure 3.**
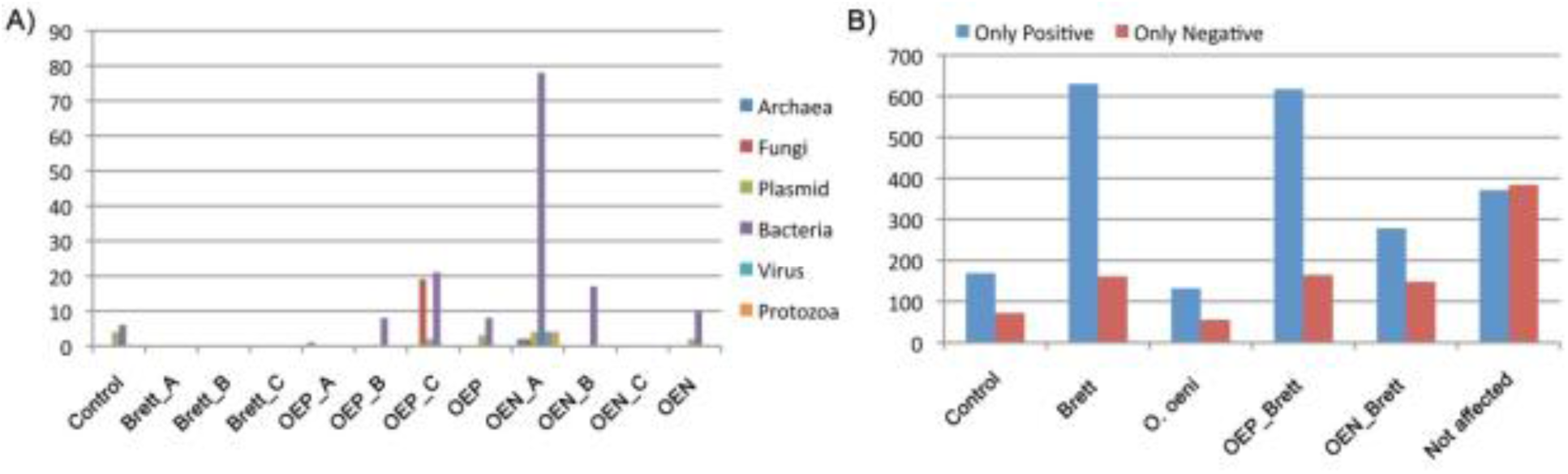
Taxonomic profiling. A) Microbial species taxonomic cores (i.e. the taxa present only in anyone of the replicates of each of the inoculation types). The general low number of identified cores highlights the low diversity in the wine microbial community, but the high number of taxa identified only in OEN_A show a high impact of the inoculation on the microbial diversity. B) Microbial abundance correlations. Correlations identified only when removing from the comparisons the values from a given inoculation type are called to be disrupted to by that inoculation type. Correlations identified regardless of removing any of the inoculation types are called “not affected correlations”.

### Abundance correlations and taxonomic distances

The number of positive and negative taxonomic abundance correlations not affected by the inoculation type (i.e. likely those that are due to the basic wine microbial community interactions) are almost the same (371 and 384, respectively) (Figure 3B). The inoculation type with the highest number of correlations disrupted by it is the addition of a *B. bruxellensis* strain (791 affected correlations), and the number of correlations affected by the OEP – *B. bruxellensis* combinations is the second highest (617, mean= 530, median= 590.5). Furthermore, the number of correlations affected by the inoculation of *O. oeni* (187), while the number of affected correlation identified by the removal of the control samples (241). Furthermore, we did not identify any correlation affected by taking into account only one specific combination of *O. oeni* and *B. bruxellensis*, but there were affected correlations identified by taking into account the three possible combinations together for each *O. oeni* strain.

As expected, the distance between the controls is the lowest than that of the other samples compared to the controls (Figure 2CD). Interestingly, the distances of the samples inoculated only with *B. bruxellensis* are similar to those of OEN_A/B/C. The distance values also show that the inoculation type with the highest difference to the controls is the OEP_A/B/C, with one of the OEP_A replicates being an out layer.

### Plasmid and fungal identifications

Similar to the bacterial community, OEP is the closest to the control in the plasmid profiling (Supplemental Figure 2A). Also, similar to the bacterial profiling, inoculation of Brett_A to OEN and OEP causes most plasmids to be in differential higher abundance than inoculation of the *O. oeni* with Brett_B or Brett_C. Interestingly, only OEP_A_10 is an out layer very distant from the pair, with about double number of identifications than the pair, although it has similar number of identified plasmids to other samples. Brett_A_3 has the fewest plasmids and OEN_A_15 has the most. However, that is not the one with highest depth, and the ones with highest depth have similar number of plasmids as the others with mean depth of sequencing. Furthermore, both Brett_A_3 and OEN_A_15 cluster together within the main cluster.

Analysis of the fungal profile clusters tightly together in most of the samples (Supplemental Figure 2B). The only three samples placed outside the main cluster with the controls are one of the replicates of the combinations of OEP with each *B. bruxellensis* strain. To evaluate whether there is an increased potential to release HCAs due to fungal activity other than that of *B. bruxellensis*, we identified in our nr gene set catalogue sequences of the cinammoyl esterase gene from fungal origin. We could only find two genes, one originally annotated as from the yeast *Pichia stipites*. However, it was not annotated as esterase, but as triacylglycerol lipase with only 31.8% identity. The other gene was originally annotated as from the yeast *Pichia pastoris*; however, it was annotated with only 30.5% identity to an uncharacterized protein.

### Profiling of functional potential

A total of 430,713 genes were predicted, with a mean of 18,726.65 per sample (Supplemental Figure 3), and a final nr gene set with all the genes from all the samples pooled was constructed containing a total of 70,991 nr genes. After filtering the low abundant ones and removing those present only in the removed OEN_23 sample for the comparative analyses, 50,604 were kept for functional annotation. A total of 41,350 of those genes were assigned a KO annotation, and we retained 40,525 nr genes after filtering out possible misannotations, accounting for 5,614 different KOs. The number of reads of the samples does not have an effect on the number of predicted genes (Pearson cor= 0.3667), thus we did not exclude samples from the functional comparative analyses based on their sequencing depth (e.g. the sample OEN_B_18, which has the highest depth of sequencing). However, we excluded sample OEN_23 because it had the lowest depth of sequencing and its functional profile was completely an out layer from the other samples, including its replicate pair (Supplemental Figure 1).

We identified a lot of functional potential variability, even within the controls (Figure 4A). OEN clusters close to OEP, which are closer to the control samples than any of the other samples. We identified variation in the functional potential between pairs of similar and different amount of sequencing. For example, the OEN_B replicates are functionally close in spite of the large difference is sequencing depth (they cluster in the same tight cluster not containing the controls). The two OEN_C replicates separate in two different tight clusters not containing the control samples and the two OEP_A replicates separate into two different clusters, one being a tight cluster containing OEP_A_9, and a loser one containing the control samples and OEP_A_10. This OEP_A pair is among the pairs most distant to each other. Furthermore, the sample with the least number of assembled nr genes was Brett_B_6 (9,184) and OEP_A_10 had the highest (41,618), although its depth of sequencing was not the lowest. The sample with the lowest depth of sequencing (OEN_23) and that with the highest (OEN_B_18) had assembled around the mean number of nr genes (14,078 and 19,651, respectively, mean= 18,219.39).

**Figure 4.**
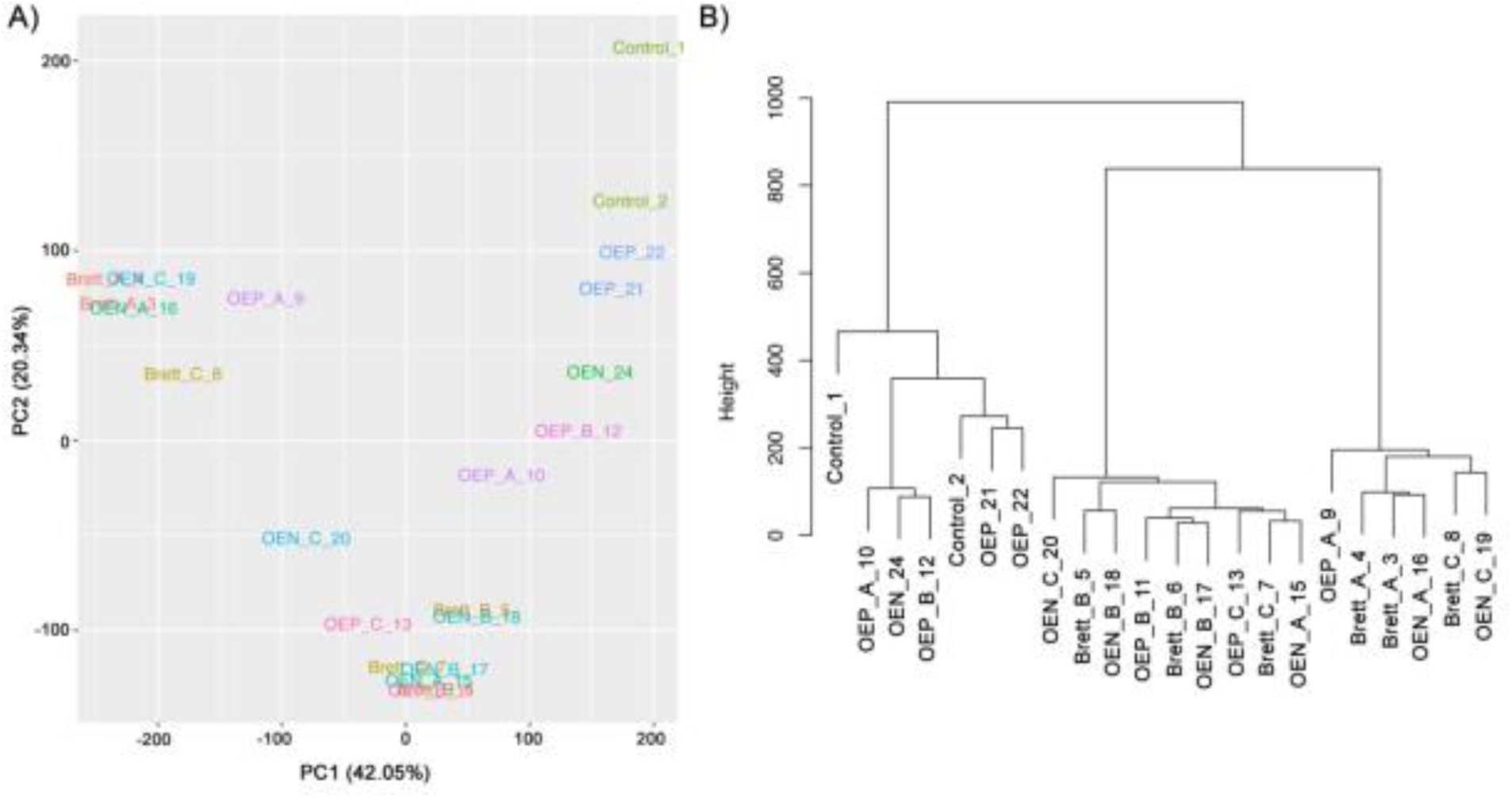
Functional profiling. A) PCA of the normalized gene mapping counts. B) Cladrogram of the Euclidean distances clustered by the ward.D method.

### Differentially abundant functions

We found that most of the differentially abundant KOs compared to the control are in lower abundance (average higher abundance= 1,221.8, median higher abundance= 1,153, average less abundance= 1,676.2, median less abundance= 1,531). However, when assigning the KOs to pathways, more pathways are affected by KOs in higher abundance (average= 36.1, median= 25) than by those in lower abundance (average= 23, median= 10). Compared to the other inoculation types, Brett_A has the highest number of differentially abundant KOs in higher abundance (2,748). Brett_A, OEP_A, OEN and OEN_C are the ones where there are more KOs in higher than in less abundance. However, OEN_A does not follow the same pattern of Brett_A and OEP_A (Figure 5A). We found that the inoculation only Brett_B and OEN_B had the highest numbers of differentially less abundant KOs compared to the control (3,481 and 3,582, respectively, average= 1,676.7, median= 1,531). OEP_B (2,757) is in the fourth place, with OEP_C in the third (3,200). *O. oeni*_B compared to Brett_B are the ones with the lowest number of KOs in differential less abundance compared to *O. oeni*_A/C, and OEN_B is the one with least KOs in higher abundance (335), although OEP_B is the one with the highest number of KOs present in higher abundance (1, 519). Brett_C has a similar number of higher and less abundant KOs (1,229 and 1,531, respectively) both around the means of higher and less of all the samples compared to the control (1,221.8 and 1,676.7, respectively). However, OEN_C is the second with the highest number of KOs present in higher abundance (2,006), and OEP_C is the third with highest number of KOs present in less abundance (3,200).

**Figure 5.**
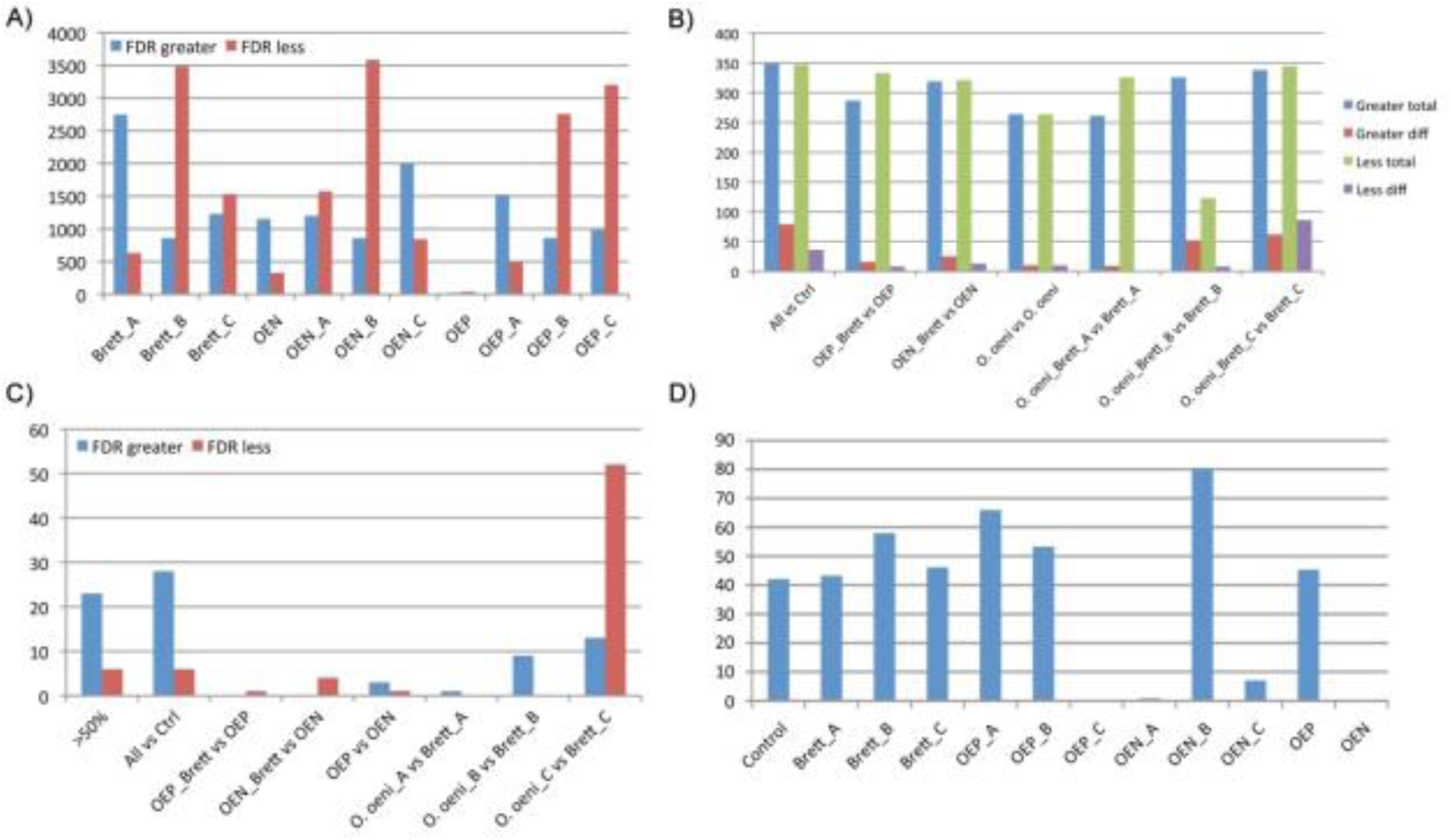
A) Differentially abundant KOs from the different inoculation types compared to the control. FDR greater denotes the KOs in statistically significant higher abundance, and FDR less are those in statistically significant less abundant. B) Pathways from the differentially abundant KOs. C) Core differentially abundant pathways from the differentially abundant KOs. D) Percentage of replicable top 5% abundant genes in each inoculation type sample pair. OEN and OEP_C did not have a replicate to consider.

OEP has the least number of higher and less abundant KOs (24 and 39, respectively) compared to the control, while inoculations with only Brett_ B and OEN_B had the highest numbers of differentially less abundant KOs compared to the control (3,481 and 3,582, respectively, average= 1,676.7, median= 1,531). Compared to OEP, OEN has more KOs present in higher abundance (956) than present in lower abundance (29). However, when looking at the pathways those KOs belong to (Figure 5B), only 10 pathways in higher and 10 in less abundance are identified, while the average is 36.14 as present in higher abundance, and 23 in lower abundance. Both replicates of inoculating only with OEP and OEN have the lowest total of differentially abundant KOs (63 and 1,477, respectively), while the inoculations with only *B. bruxellensis* have larger numbers of differentially abundant KOs. When analysing the pathways differentially abundant in only a given inoculation type derived from differentially abundant KOs, the comparisons of both OEP_C and OEN_C have the highest number of differentially less abundant pathways (52) (Figure 5C), with most of those differentially abundant KOs coming from OEP_C (2,068, while OEN_C has 593).

### Core and top abundant genes

Out of the 40,525 annotated filtered nr genes, only 867 are present in all the compared samples. From all the *B. bruxellensis* and *O. oeni* – *B. bruxellensis* combinations, Brett_A has the highest number of genes in its core (i.e. present in the two replicates of the inoculation type), with only 15. Inoculations of Brett_A alone have only 1 core gene (annotated as coming from *Candida glabrata*), while the other two inoculations of Brett_B and Brett_C have zero genes in the core. The *O. oeni*_B inoculations do not have genes in the core either, while *O. oeni*_C has only 9. OEN has only 16 core genes, much less core genes than OEP (329), but still in the third place compared to the other inoculation types. It is interesting that although OEP is the closest to the control, it is the second with the highest number of genes in its core (329, median= 2, mean= 67.15). As expected, the control samples are the ones with most genes in its core (515).

A total of 1,582 different nr genes were found among the top abundant genes (*P* < 0.05) of all the inoculation types. The percentage of top abundant genes present in both replicates of each type of inoculation is in average 53.49% (median of 49.64%), highlighting the variability in the wine microbial community. OEN_A has the lowest number of replicable genes present in top abundance in both replicates (Figure 5D) (only 5 genes), while both replicates have around the mean number of top abundant genes (580 and 646, mean=627.67). Notably, OEN_B has the highest percentage of replicable top abundant genes from the replicates (80.2%, 355, each replicate has 438 and 447, which are below the mean and median).

### HCA derivatives

To evaluate whether an increase in the release of HCA has the potential to produce more off-flavour compounds, we looked for genes in our nr gene set catalogue involved in the processing of HCAs. We identified a gene from *Erwinia gerundensis* (a cosmopolitan epiphyte) with 35.294% identity annotated with the KO K13727 (phenolic acid decarboxylase), which has decarboxylation activity on HCAs ferulic, p-coumaric and caffeic acids. This same gene had a second putative annotation with 73.864% identity to an unannotated protein from *Nectria haematococca* (*Fusarium solani* subsp. pisi), which is a fungal plant pathogen. This gene was also found to be significantly more abundant in Brett_A compared to Brett_B (FDR= 4.16e^−10^), OEP_A (FDR= 0.0138), OEN_A (FDR= 6.13e^−5^), and control (FDR= 2.91e^−8^). It was also more abundant in Brett_C than in OEP_C (FDR= 0.0027) and the control (FDR= 4.54e^−7^), in OEN_C than in OEN (FDR= 1.27e^−11^) and in control (FDR= 3.22e^−11^), and in OEP_A than in OEP (FDR=0.0014) and control (FDR=0.0012). We also identified another KO related to the carboxylation of HCAs among our assembled genes; K20039 (ferulic acid decarboxylase 1-like) from *S. cerevisiae*, which was in significant higher abundance in OEP_A compared to Brett_A (FDR= 0.00014) and in OEP_B compared to Brett_B (FDR= 0.02). Also, some genes with this KO were identified as less abundant in OEP_B and OEP_C compared to OEP (FDR= 0.01 and FDR= 3.82e^−9^, respectively), and the control (FDR= 0.013, FDR= 2.10e^−9^, respectively). Other genes annotated as ferulic acid decarboxylase 1-like were in significant less abundance in OEN_A/B/C compared to OEN (FDR= 2.1e^−10^, FDR= 5.25e^−8^, FDR= 4.17e^−8^, respectively) and the control (FDR= 1.19e^−7^, FDR= 1.88e^−5^, FDR= 8.92e^−6^), and less abundant in the Brett_A/B/C compared to the control (FDR= 1.99e^−6^, FDR= 3.55e^−7^, FDR= 1.23e^−8^, respectively).

## Discussion

### Microbial taxonomic and functional potential profiling

#### General wine-related identifications

Among the taxonomic identifications, we found expected bacteria derived from soil and plants, such as *Xanthomonas alfalfa*, *Dyella japonica* and *Micrococcus luteus* (found on the surface of table grapes). We also identified wine spoilage bacteria, such as *Aeromonas hydrophila* in various samples. In regards to the number of bacterial identifications, it is interesting to note that we found *Bodo saltans* (*Pleuromonas jaculans*) in top abundance in OEP_B_12 (at 308.2x coverage), OEN_24 (126.57x), both OEN_C replicates, OEN_A_16, and Brett_A_3 (mean coverage of 21.57x). It is a free-living nonparasitic protozoan which feeds upon bacteria which can be found in freshwater and marine environments. These samples that have it in top abundance also have less bacterial identifications compared to the rest or compared to their respective pair. However, the causal relationship cannot be identified, although it deserves further study.

Among the top abundant bacteria identified in OEP combined with a *B. bruxellensis* strain, we found several LAB that are examples of bacteria being promoted as an effect of the use of a specific *O. oeni* strain with *B. bruxellensis*. For instance, we identified *Lactobacillus collinoides*, a LAB found in cider, in one of the replicates of each OEP_A/B/C combination in higher abundance compared to the respective *B. bruxellensis* strain and to OEP and the controls. In similar differential abundance pattern, we found among the top abundant bacteria *Lactobacillus crustorum*, isolated from two traditional Belgian wheat sourdoughs, *Lactobacillus herbarum*, a species related to *Lactobacillus plantarum*, *Lactobacillus oeni*, LAB isolated from wine, and *Lactobacillus paucivorans*, isolated from a brewery environment. Among the top abundant bacteria identified in higher differential abundance in each OEP_A/B/C compared to the respective *B. bruxellensis* strain, to OEP and the controls, we identified *Lactobacillus nagelii,* isolated from a partially fermented wine, and *Lactobacillus parafarraginis,* a heterofermentative lactobacilli isolated from a compost of distilled shochu residue.

Among the fungi in differentially higher abundance in OEN_A when compared to Brett_A, OEN and the control, and in OEN_B/C when compared to OEN, we identified *Talaromyces stipitatus*. This fungus contains genes with high identity to those needed for the biosynthesis of the red pigment monascorubrin by the phylogenetic relative fungi *Talaromyces marneffei*. Interestingly, we also identified *T. marneffei* differentially abundant only in in OEP_C when compared to Brett_C, OEP and control. As expected, we also identified other grape-related fungi, such as the plant pathogens *Verticillium dahlia* and *Verticillium longisporum, Mucor ambiguus*, present in soil and plants, and the wine common yeast *Saccharomyces cerevisiae* and *Lachancea kluyveri* (found in the core of OEP_C).

In regards to AF, we identified in various samples the fungi *Mucor indicus*, previously isolated from the traditional fermented Indonesian food tempeh, and with the capability of producing ethanol is comparable with that of *S. cerevisiae*. Also, in one OEP_C sample we identified *Candida sorboxylosa*, an ethanol producing and tolerant yeast species from fruits for production of bio-ethanol that is common to the winery environment.

#### Microbial community identification comparisons

We identified no malic acid on day 114 in the samples not inoculated with our *O. oeni* culture strains (Figure 1), suggesting that the found high coverage of *O. oeni* in the samples not inoculated with them (one replicate of Brett_A and Brett_C, and the two Brett_B replicates) derives from *O. oeni* present in the native grape flora. From the three tested *B. bruxellensis* strains, Brett_B seems to be unable to grow in these wines, because when inoculated alone it was identified at only 0.4x, and when inoculated with *O. oeni*, it was identified in very low abundance (Table 1, Supplemental File 1). The difference in the genome depths of coverage in the replicates of the *O. oeni* and *B. bruxellensis* strains inoculated together suggest that *B. bruxellensis* sometimes grows well and sometimes not, depending on certain unidentified conditions. Such inability to predict with precision the activity of a spoilage yeast and its effect on the entire microbial community highlights the importance in wine-making of inoculating with sufficient numbers of strong and viable yeast and bacteria to ensure the presence of the desired microbial community (Gerbaux et al. 2009). In the identification of bacterial species from plasmids, all the inoculation types seem to have an effect on the community compared to the controls, while on the bacterial community there was an effect only for certain inoculation types. This suggests that the plasmids presence is inherently variable, although to a low extent. The fact that OEP_A_10 has similar number of identified plasmids to other samples in spite of being an out layer very distant from the pair (with about double number of identifications than the pair) suggests that the difference in the plasmid profile resides on the bacterial community (the plasmids hosts). Also, the differences in the number of plasmids identified in the samples compared to their depth of sequencing, suggests that the number of identifications is not mainly due to depth of sequencing.

Interestingly, while the number of reads of the samples has a moderate influence on the number of identified taxa (Pearson cor=0.6832), it does not have an effect on the number of identified genes (Pearson cor= 0.3667). Furthermore, the number of identified taxa has a moderate correlation to the number of identified genes (Pearson cor= 0.6854), and the number of nr genes directly correlates with the number of genes in all the samples (Pearson cor= 0.997) (Supplemental Figure 3). This suggests that the functional potential space of the wine microbiome is more defined than the taxonomic profile. However, we also identified functional potential variation between pairs of similar and different amount of sequencing, suggesting that the observed variability in the functional potential is not due to differences in depth of sequencing. Interestingly, the variability is such that even the controls do not cluster tightly together as in the taxonomic PCAs. But, similarly to the impact of *B. bruxellensis* in the taxonomic identifications, most of the impact when *O. oeni* is inoculated occurs when combining it with a *B. bruxellensis*, as OEN and OEP are the closest to the control samples. These results suggest that the main changes in the wine microbial community occur when *B. bruxellensis* is present.

In contrast to the taxonomic profiling, where most of the taxa of the different inoculation types compared to the control was in statistically significant higher abundance, at the functional level most of the differentially abundant KOs compared to the control are in less abundance. However, the assignation of the KOs to pathways showed that more different pathways are affected by KOs in higher abundance than by those in less abundance. This suggests that there is a need of a minimum set of present pathways required by the wine microbiome to tribe in that particular system; such minimum set can be perturbed by changes in the component KOs, however such changes do not disrupt the entire pathway. Elimination of the presence of a KO (i.e. differentially less abundant KO) is more disruptive to a pathway than a KO being present in higher abundance.

The fact that only the control and OEP have a large number of genes as core, but that the OEP and the control samples are functionally the closest types of inoculation, suggests that the main effect on the functional profiles of the different inoculations is not in the integration of new functions, but in changes in their abundance.

#### O. oeni *-* B. bruxellensis *strain specific dependent effect*

We observed that in general, the samples inoculated with *B. bruxellensis* have less bacterial identifications (mean 71, average all samples 152.4, median all samples 129) than the ones inoculated with an *O. oeni* strain (Supplemental File 1). This suggestion is supported by the correlation abundance results. If the effect of the addition of a *B. bruxellensis* strain is the elimination of several taxa, it would be expected that most of the affected correlation were those that used to be positive. Effectively, 79.6% of the affected correlations were positive. The fact that both replicates of inoculating only with OEP and OEN cluster together with the controls and have the lowest total of differentially abundant KOs (63 and 1477, respectively), while the inoculations with only *B. bruxellensis* strains have larger numbers of differentially abundant KOs, suggests that most of the functional potential impact is given by the *B. bruxellensis* strain than by the *O. oeni* strain.

Also, the clustering patterns on the PCA of the taxonomic identifications suggest that the combination of *O. oeni* and *B. bruxellensis* has an impact on the bacterial composition depending on the strains being combined, both between and within inoculation types. It also suggests that in some instances the presence of more of low abundant bacteria and difference in abundance of the same bacteria accounts for the variation within the same inoculation type, rather than a radical change in the bacterial composition.

The observation that *O. oeni* and *B. bruxellensis* have an impact on the bacterial composition depending on the specific strains being combined, both between and within inoculation types, is supported by the abundance correlation analyses. Correlations were not affected when taking into account only one specific combination of *O. oeni* and *B. bruxellensis*, but when taking into account the three possible combinations together for each *O. oeni* strain. In other words, the effect of OEP_A is not the same as that of OEP_B and OEP_C, and also that of OEN_A is not the same as OEN_B and OEN_C. Also, when looking at the functional potential evaluation, we found that compared to the other inoculation types, Brett_A and OEP_A are among those with the highest number of differentially higher abundant KOs. However, OEN_A does not follow the same pattern of Brett_A and OEP_A. This further supports the suggestion that the functional impact of Brett_A depends on the *O. oeni* it interacts with (Figure 5A).

Interestingly, in spite of the observation that OEN_B is the one with least KOs in greater abundance while OEP_B is the one with the highest number of KOs present in higher abundance, the inoculations of only Brett_B, OEN_B, and OEP_B are among the samples with the highest numbers of differentially less abundant KOs compared to the control. This suggests that Brett_B reduces the functional potential of the system, regardless of the *O. oeni* it is inoculated with. On the other hand, Brett_C has a similar number of higher and less abundant KOs compared to the control. However, OEN_C is the second with the highest number of KOs present in higher abundance and OEP_C is the third with highest number of KOs present in less abundance. This suggests that the effect of Brett_C on the functional potential depends on the *O. oeni* strain.

Notably, although OEN_A and OEN_C have the least number of replicable top abundant genes, the OEN_B samples have the highest percentage of replicable top abundant genes, suggesting that the OEN_B functional profile is more replicable and less variable than that of the other inoculation types, again highlighting the different impact in the microbial communities that *O. oeni* strains have depending on the *B. bruxellensis* strain present.

The observation that the inoculations of Brett_A together with both OEN and OEP have low numbers of differentially less abundant bacteria suggests that the inoculation of Brett_A with *O. oeni* seems to impact the community by allowing the growth of more bacterial taxa instead of by repressing their growth. Although the large variability within the OEP_A inoculation type is likely due to the ability of the Brett_A to grow poorly or successfully with OEP, this same effect is not observed in OEN_A, bec ause in OEN_A one replicate grew more than the other, but the core of the OEN_A inoculation type is the largest (Figure 3A). Among them, we found *Lactobacillus paracollinoides*, isolated from brewery environments, *Lactobacillus pentosus,* the most prevalent lactic acid bacterium in Spanish-style green olive fermentations, and *Staphylococcus equorum*, frequently isolated from fermented food products and contributing to the formation of aroma compounds during ripening, especially in cheeses and sausages.

### O. oeni strain *specific effect*

OEN had the lowest number of bacteria present in higher abundance compared to the control, and OEP is the closest to the control and is among the inoculation types with least bacteria present in higher abundance. This suggests that the inoculation type that affects the least the wine microbial profile (after 6 months of inoculation) is that of a single *O. oeni* strain. This is supported by the microbial abundance correlation analyses, where the number of correlations affected by the inoculation of *O. oeni* is close to the number of affected correlation identified by the removal of the control samples. Furthermore, OEP has the least number of higher and less abundant KOs compared to the control, and its functional potential diversity clusters together with the controls (Figures 4B, 5A), suggesting that OEP causes the least change in the functional potential compared to the other inoculation types. Compared to OEP, OEN has more KOs present in higher abundance than present in less abundance, however, when looking at the pathways those KOs belong to (Figure 5B), only 10 pathways in higher and 10 in less abundance are identified. These observations suggest that the degree of functional impact of those *O. oeni* strains alone is similar.

We observed low numbers of replicable genes present in the 5% top abundant genes in both replicates of OEN_A (Figure 5D), although they have around the mean number of top abundant genes. This is likely due to the difference in abundance of *O. oeni*; one of the replicates had *O. oeni* in high abundance, while the other had the *O. oeni* in very low abundance and the *B. bruxellensis* in moderate abundance.

### The effect of the abundance of B. bruxellensis

Interestingly, in the viral profiling, in OEN_A we identified in higher abundance compared to the control two viruses against fungi, *Phytophthora infestans* RNA virus 1 and *Saccharomyces cerevisiae* killer virus M1 (also in higher abundance in OEN_B). It could be that these viruses contribute to a reduction of the fungal diversity in this OEN – *B. bruxellensis* combinations compared to those combined with OEP, where fungal diversity seems to be increased. However, this would need further experimental validation.

Notably, when checking the pathways differentially abundant in only a given inoculation type derived from differentially abundant KOs, the comparisons of *O. oeni*_C versus Brett_C have the highest number of differentially less abundant pathways (Figure 5C), with most of those differentially abundant KOs coming from OEP_C. This suggests that OEP combined with a successfully growing Brett_C causes a large impact on the functional potential of the wine microbial community. Among the differentially abundant pathways present in less abundance in the comparisons of *O. oeni*_C versus Brett_C is the regulation of mitophagy in yeast, with all the KOs of the pathway in less abundance in OEP_C, possibly suggesting there is less potential of regulation of the fungal taxonomic profile in OEP_C. This is interesting, as the fungal profile in OEP_C is the second with the highest number of differentially more abundant fungal species when compared to the control (61, while the mean is 30.54 and median is 14).

The observation that OEP alone is closest to the control in the taxonomic profiling, and that it has a strong effect on the bacterial profile when *B. bruxellensis* is in high abundance is notable, given that HCAs have been shown to inhibit the growth of many microorganisms (Kheir et al. 2013).

Furthermore, there is evidence that the concentration of 4-ethylphenol and 4-ethylguaiacol is lower when malolactic bacteria are present before exposure to *Brettanomyces*, so that it has been suggested to inoculate with commercially available strains as the time needed for spontaneous MLF is unpredictable giving prolonged risk for exposure of *B. bruxellensis* (Nielsen & Richelieu 1999).

### Flavour potential

#### Taxonomic and functional potential identifications

In regards to taxonomic identifications related to flavour formation in wine, we found only in the OEP_C sample the bacterium *Lactobacillus diolivorans* (176 mapping reads), which degrades 1,2-propanediol, a compound that is nearly odourless but that possesses a faintly sweet taste. Also, only in this sample we found the fungi *Clavispora lusitaniae* (596 mapping reads), which has been found to produce a good balance between concentrations of ethyl acetate (sweet smell) and high alcohols. In regards to potential functions, the β-glucosidase activity is involved in the hydrolysis of several important compounds for the development of varietal wine flavour profiles, and microbial β-glucosidases have been used for the enhancement of wine aroma. Importantly, glucosydases not encoded by *S. cerevisiae* have been shown to impact the flavour compounds profile in wine (Rosi et al. 1994). In our nr gene set we identified 53 genes annotated as KO K05349 (bglX; beta-glucosidase) from many different bacteria and non-*Saccharomyces* yeast. One of them is in top abundance in various samples and is annotated as coming from *O. oeni*. However, sensorial evaluation is required to assess the impact in the wine flavour due to these identified genes.

### MLF

To evaluate whether the presence of *B. bruxellensis* affects the occurrence of MLF, and thus its effect in reducing the acidity of the wine, we looked for genes annotated as malate dehydrogenase (*mdh*) in our nr gene set. We identified *mdh* in differential abundance in various comparisons and coming from various species, including *O. oeni*. As expected, the two genes annotated as *mdh* and D-lactate dehydrogenase from *O. oeni* are in the top abundant in all the samples inoculated with *O. oeni* strains and *O. oeni* combined with a *B. bruxellensis* strain, and also in Brett_B/C. Thus, the presence of the analysed *B. bruxellensis* strains does not affect the MLF activity of the evaluated *O. oeni* strains.

### HCA production

The cinnamoyl esterase activity can also be present in different fungi, suggesting that under certain unidentified conditions, the variability of fungi promoted by the combination of OEP with a well growing *B. bruxellensis* could possess this esterase activity and contribute to the increase in the production of HCAs. However, we could only find two yeast genes with inconclusive functional annotations. Thus, it cannot be concluded that there is a higher production of HCA due to the potential activity of other fungi, further experimental functional characterization should be performed on these identifications to validate whether they confer the cinammoyl esterase activity. The observations from the abundances of the phenolic acid decarboxylase and ferulic acid decarboxylase 1-like genes also suggest that the presence of genes with putative decarboxylase activity on HCAs is not dependent on the presence of a specific *O. oeni* strain with or without the esterase activity, but on the *B. bruxellensis* strain, because it is also in the control samples and because Brett_A has higher abundance of phenolic acid decarboxylase than OEP_A.

## Conclusions

In this study, we characterized the impact on the microbial community of a Cabernet Sauvignon wine six months post-inoculation of two different strains of *O. oeni* (with and without the cinnamoyl esterase activity) and three *B. bruxellensis* strains, alone and in combination. We found that the impact in the taxonomic profile and functional potential of the microbiome due to the *O. oeni* – *B. bruxellensis* combinations depends on *i*) the specific *O. oeni* and *B. bruxellensis* strains being combined, and *ii*) the abundance reached by the inoculants, which depends on certain unidentified conditions. Analysis of the functional potential of the system identified that changes in the abundance of the genes is the general effect of the inoculations, not integration of new functions. OEP maintains the stability of the most abundant functions in the system in spite of the addition of the *B. bruxellensis*. The functional potential for the HcD activity is dependent on the *B. bruxellensis* strain; however, the control samples also have this potential, not derived from the inoculated *B. bruxellensis* nor from the *O. oeni* strains. Furthermore, it was not possible to identify non-Brettanomyces fungal potential to produce HCAs as a result from a particular inoculation type. Finally, the HCAs post processing into off-flavor compounds is not dependent on the *O. oeni* but on the *B. bruxellensis* strain and other microbes, likely derived from the indigenous grape flora. This study proves the usefulness of metagenomic analysis in obtaining a deeper insight into the general microbial profile characteristics and the impact of specific inoculants, not only in the taxonomy, but also in the functional potential of the system. However, experimental validation will be necessary in future studies to obtain a detailed knowledge of the specific mechanisms of the interactions identified with metagenomic analyses. Also, sensorial analysis would be necessary to evaluate the impact on the flavor profile of the wines produced by the potential unveiled by the metagenomic results.

## Acknowledgements

MLZM, MAK and LP thank the Danish National Advanced Technology Foundation (Højteknologifonden) 080-2012-3-Food genomics and Innovation Fund Denmark case number 6150-00033A FoodTranscriptomics for funding the research. The authors declare no conflict of interests. We thank the Danish National High-Throughput DNA Sequencing Centre for the generation of the sequencing data. We gratefully acknowledge the Danish National Supercomputer for Life Sciences – Computerome (computerome.dtu.dk) for the computational resources to perform the sequence analyses. The sequencing data reported in this paper will be archived in a public database (ID added upon paper acceptance).

Short version of title: *O. oeni* - *B. bruxellensis* impact on wine microbiome

## Literature Cited

Barata A et al. 2008. Ascomycetous yeast species recovered from grapes damaged by honeydew and sour rot. Journal of applied microbiology 104(4):1182–91.

Barata A, Malfeito-Ferreira M and Loureiro V. 2012. The microbial ecology of wine grape berries. International Journal of Food Microbiology 153(3):243–259.

Bokulich NA et al. 2014. Microbial biogeography of wine grapes is conditioned by cultivar, vintage, and climate. Proceedings of the National Academy of Sciences of the United States of America 111(1):E139–48.

Bouzanquet Q et al. 2012. A Novel Glutathione-Hydroxycinnamic Acid Product Generated in Oxidative Wine Conditions. Journal of Agricultural and Food Chemistry 60(49):12186–12195.

Branco P et al. 2014. Identification of novel GAPDH-derived antimicrobial peptides secreted by Saccharomyces cerevisiae and involved in wine microbial interactions. Applied Microbiology and Biotechnology 98(2):843–853.

Cabrita MJ et al. 2008. Impact of malolactic fermentation on low molecular weight phenolic compounds. Talanta 74(5):1281–1286.

Edgar RC. 2010. Search and clustering orders of magnitude faster than BLAST. Bioinformatics (Oxford, England) 26(19):2460–1.

De Filippis F, Parente E and Ercolini D. 2017. Metagenomics insights into food fermentations. Microbial Biotechnology 10(1):91–102.

Gerbaux V et al. 2009. Influence of Inoculation with Malolactic Bacteria on Volatile Phenols in Wines. American Journal of Enology and Viticulture 60(2):233–235.

Harris RS. 2007. Improved Pairwise Alignment of Genomic DNA. ProQuest.

Hernández T et al. 2006. Phenolic compounds in red wine subjected to industrial malolactic fermentation and ageing on lees. Analytica Chimica Acta 563(1-2):116–125.

Hixson JL et al. 2012. Hydroxycinnamic Acid Ethyl Esters as Precursors to Ethylphenols in Wine. Journal of Agricultural and Food Chemistry 60(9):2293–2298.

Hyatt D et al. 2010. Prodigal: prokaryotic gene recognition and translation initiation site identification. BMC bioinformatics 11:119.

Kheir J et al. 2013. Impact of volatile phenols and their precursors on wine quality and control measures of Brettanomyces/Dekkera yeasts. European Food Research and Technology 237(5):655–671.

Kikuzaki H et al. 2002. Antioxidant properties of ferulic acid and its related compounds. Journal of agricultural and food chemistry 50(7):2161–8.

Leff JW et al. 2013. Bacterial Communities Associated with the Surfaces of Fresh Fruits and Vegetables. PLoS ONE 8(3):e59310.

Li H and Durbin R. 2009. Fast and accurate short read alignment with Burrows-Wheeler transform. Bioinformatics (Oxford, England) 25(14):1754–60.

Liu SQ. 2002. A review: malolactic fermentation in wine -- beyond deacidification. Journal of applied microbiology 92(4):589–601.

Liu Y et al. 2017. Wine microbiome: A dynamic world of microbial interactions. Critical Reviews in Food Science and Nutrition 57(4):856–873.

Lonvaud-Funel A. 1999. Lactic acid bacteria in the quality improvement and depreciation of wine. Antonie van Leeuwenhoek 76(1-4):317–31.

Madsen MG et al. 2016. Influence of Oenococcus oeni and Brettanomyces bruxellensis on Hydroxycinnamic Acids and Volatile Phenols of Aged Wine. American Journal of Enology and Viticulture doi: 10.5344/ajev.2016.16015.

Martin M. 2011. Cutadapt removes adapter sequences from high-throughput sequencing reads. EMBnet.journal 17(10).

Meyer M and Kircher M. 2010. Illumina sequencing library preparation for highly multiplexed target capture and sequencing. Cold Spring Harbor protocols 2010(6):pdb.prot5448.

Nagel CW and Wulf LW. 1979. Changes in the Anthocyanins, Flavonoids and Hydroxycinnamic Acid Esters during Fermentation and Aging of Merlot and Cabernet Sauvignon. American Journal of Enology and Viticulture 30(2):111–116.

Nielsen JC and Richelieu M. 1999. Control of flavor development in wine during and after malolactic fermentation by Oenococcus oeni. Applied and environmental microbiology 65(2):740–5.

Ou S and Kwok KC. 2004. Ferulic acid: pharmaceutical functions, preparation and applications in foods. Journal of the Science of Food and Agriculture 84(11):1261–1269.

Peng Y et al. 2012. IDBA-UD: a de novo assembler for single-cell and metagenomic sequencing data with highly uneven depth. Bioinformatics (Oxford, England) 28(11):1420–8.

Petersen TN et al. 2017. MGmapper: Reference based mapping and taxonomy annotation of metagenomics sequence reads. PLOS ONE 12(5):p.e0176469.

Pina C et al. 2004. Ethanol tolerance of five non-Saccharomyces wine yeasts in comparison with a strain of Saccharomyces cerevisiae influence of different culture conditions. Food Microbiology 21(4):439–447.

Rosi I, Vinella M and Domizio P. 1994. Characterization of beta-glucosidase activity in yeasts of oenological origin. The Journal of applied bacteriology 77(5):519–27.

Rumbold K et al. 2003. Purification and properties of a feruloyl esterase involved in lignocellulose degradation by Aureobasidium pullulans. Applied and environmental microbiology 69(9):5622–6.

Sieuwerts S et al. 2008. Unraveling microbial interactions in food fermentations: from classical to genomics approaches. Applied and environmental microbiology 74(16):4997–5007.

Sipiczki M. 2006. Metschnikowia strains isolated from botrytized grapes antagonize fungal and bacterial growth by iron depletion. Applied and environmental microbiology 72(10):6716–24.

Suarez R et al. 2007. The production of ethylphenols in wine by yeasts of the genera Brettanomyces and Dekkera: A review. Food Chemistry 102(1):10–21.

